# UVC Irradiation Induces Time-Dependent Proteomic Remodeling in Geodermatophilus obscurus G20

**DOI:** 10.64898/2026.11.04.764414

**Authors:** Brandon Anselmo Saiz, Seifeddine Ben Tekaya, Naomi Ward, Franco Basile

## Abstract

*Geodermatophilus obscurus* G20 is a multi-resistant extremophile observed to survive significant dosages of ultraviolet (UV) radiation. Despite previous work having measured *G. obscurus* G20’s proteome in relation to its genome sequence, its proteomic response to stressors has yet to be explored. This study utilizes bottom-up proteomics with data independent acquisition to derive comprehensive proteomic measurements in both healthy and two UV-stressed conditions induced through 254 nm UV radiation administered at a median lethal dose. The first, an acute response treatment group, was recovered and flash-frozen immediately after UV radiation. The second, a recovery response treatment group, was allowed to incubate for 1 hour following UV radiation. Gene enrichment analysis revealed an increase in NADH dehydrogenase activity among both stressed treatment groups, however the recovery response treatment group experienced additional changes including reduced peptidoglycan synthesis and increased translational activity. Results suggest that the repair process undergone by *G. obscurus* G20 persists well after initial UV exposure and is likely to be inducing noteworthy energetic demands to facilitate UV induced genomic repair. Proteomic analyses revealed patterns that advance our mechanistic and temporal understanding of the *G. obscurus* metabolic response to UV irradiation.

## Introduction

As anthropogenic changes to the atmosphere have highlighted the protective role of the stratospheric ozone layer, understanding how organisms survive ultraviolet (UV) radiation has become increasingly important (Bernhard et al. 2023). Ultraviolet radiation is a ubiquitous abiotic stressor that threatens cellular integrity across all domains of life. Among its many biological effects, DNA is highly susceptible to UV radiation, where absorption of UV energy promotes the formation of covalent bonds between adjacent pyrimidine bases (Nieto Moreno et al. 2023). This modification within the DNA distorts its helix structure and disrupts DNA-binding protein activity, such as transcription factors and replication machinery, in the effected region (Tommasi et al. 1996; Han et al. 2016; Walsh et al. 2026). Although nucleotide excision repair (NER) and related repair pathways correct many UV-induced lesions, these repairs are error prone, and present an increased rate of mutation following their involvement (Douki et al. 2017; Nieto Moreno et al. 2023). Consequently, microorganisms inhabiting highly irradiated environments require efficient mechanisms to repair and tolerate UV-induced DNA damage.

Among radiation-resistant bacteria, *Geodermatophilus obscurus* G20 has attracted considerable interest due to its exceptional tolerance to ultraviolet irradiation, making it an excellent model for investigating the molecular basis of microbial radiation resistance (Gtari et al. 2012; Sghaier et al. 2016). *Geodermatophilus obscurus* G20 is a gram positive member of the phylum Actinomycetota, that was first isolated from the sun-exposed soils of the Amargosa Desert in Nevada, USA (Luedemann 1968). Since then, members of the genus *Geodermatophilus* have been repeatedly isolated from a wide range of harsh terrestrial environments, including arid desert soils, exposed rock surfaces, and stone monuments subjected to intense solar irradiation (del Carmen Montero-Calasanz et al. 2013; Montero-Calasanz et al. 2013a, b; Hezbri et al. 2015, 2016; Ben Tekaya et al. 2024). The extent of this resistance was quantified by Sghaier et al. (2016), who reported a D_10_ dose of 3,500 kJm^-2^ under 254 nm UV irradiation, approximately 583 times higher than its phylogenetic relative *Blastococcus saxobsidens*. The authors further demonstrated resistance to the DNA cross-linking agent mitomycin C, suggesting that the observed UV tolerance could not be explained solely by carotenoid-mediated UV shielding. Here, we investigated the proteomic response of *Geodermatophilus obscurus* G20 to ultraviolet irradiation by exposing cells to a median lethal dose (LD_50_) of 254-nm UV light. Proteomes of the exposed cells were measured immediately following irradiation (acute response) and after a 1-hour recovery period.These conditions were selected based on previous work examining the UV response of *Deinococcus radiodurans* (Patel et al. 2009). Detected proteins were quantified using bottom-up label-free quantitative (LFQ) proteomics to identify the molecular pathways underlying the acute and early recovery responses to UV stress.

## Materials and Methods

### LD_50_ Measurement

*G. obscurus* was streaked onto solid Luedmann plates and irradiated via Stratalinker 254 nm UV at various time intervals with 3-6 biological replicates per interval. Subsequent colony growth indicated the resultant colony forming unit (CFU) counts observed at each interval. An exponential line of best fit was used to infer the time at which the median lethal dose to exist.

### *G. obscurus* Liquid Cultures

Colonies from freshly streaked Luedemann agar plates were used to inoculate 5 mL starter cultures in Luedemann medium. Following incubation, the starter cultures were homogenized under sterile conditions using a sterile pestle and then repeatedly passed through a sterile syringe fitted with a 26G needle, following a protocol adapted from Gtari et al. (2012). A 1 mL aliquot of the homogenized starter culture was used to inoculate 100 mL Luedemann medium in a 250 mL Erlenmeyer flask, corresponding to a 1% (v/v) inoculum. Cultures were incubated at 30 °C for approximately 12 days.

### UV Dosage & Sample Collection

Following growth in liquid culture, contents from the Erlenmeyer flasks were transferred to 50 mL falcon tubes and centrifuged for 10 minutes at 4,696 RPM using a Thermo Scientific Sorvall ST 40 Benchtop Centrifuge. Liquid supernatant was then discarded while the cell pellet at the bottom was retained. This was repeated to transfer all cell mass from the Erlenmeyer flask into the falcon tube, and to discard the original Luedmann media the cultures were grown in. Samples were then washed in 45 mL of 0.9% NaCl solution, and then washed again in 25 mL NaCl solution before centrifugation and discarding the NaCl supernatant. At this step, non-irradiated control samples (n = 9) were submerged in liquid nitrogen then stored at -80 °C for future proteomic extraction. Treatment samples were then suspended in 0.9% NaCl solution, and vortexed to resuspend and break apart the cell pellet.

Preceding UV exposure, treatment samples were poured into empty 150 x 15 mm (diameter x height) petri dishes. Once transferred, samples were placed into a Stratagene Stratalinker 1800 and irradiated using “time mode” for 13 seconds, corresponding to an estimated dose of approximately 39 mJ/cm² based on the manufacturer-specified irradiance of ∼3000 µW/cm² (Stratalinker 2400 UV Crosslinker, Stratagene, La Jolla, CA, USA) before being removed from the Stratalinker. Following irradiation, cell mass was transferred along with surrounding NaCl solution via pipette into a new 50 mL falcon tube. Samples belonging to the acute treatment group (n = 9) were then centrifuged for 10 minutes at 4,696 rpm, their NaCl solution was discarded, and samples were submerged in liquid nitrogen to be flash-frozen before subsequent storage at -80 °C. Samples belonging to the recovery treatment group (n = 9) were allowed to incubate at room temperature for 1 hour within falcon tubes before undergoing the same centrifugation and flash-freezing process previously described. It is important to note that recovery treatment group samples were irradiated first, and the acute treatment group samples were suspended in NaCl solution while the recovery treatment group incubated, resulting in both groups being suspended in NaCl solution for equal amounts of time.

### Protein Extraction and Sample Preparation for Proteomic Analyses

Frozen samples were resuspended in 1 mL lysis buffer (4% SDS, 100 mM Tris-HCl, pH 8.0, 10 mM dithiothreitol) and sonicated using a VWR Ultrasonic Cleaner (97043-964) at 80% amplitude for 2.5 min with 10 s on/off cycles (5 min total). Samples were subsequently boiled for 5 min, and proteins were precipitated overnight at −20 °C in 80% acetone. Following centrifugation at 5,000 rpm, protein pellets were washed twice with 100% acetone and air-dried.

Dried pellets were resuspended in 500 μL digestion buffer (8 M urea, 100 mM Tris-HCl, pH 8.0, 5 mM dithiothreitol), briefly sonicated on ice (2 s at 30% intensity), and incubated for 30 min at room temperature. Proteins were alkylated with 20 mM iodoacetamide for 15 min in the dark. Samples were then diluted with 2 mL of 100 mM Tris-HCl (pH 8.0) and digested overnight at 37 °C with 5 μg sequencing-grade modified trypsin (Promega, V5111).

Following digestion, samples were acidified to 1% formic acid and peptides were purified using C18 solid-phase extraction cartridges (Waters Sep-Pak, WAT054955). Cartridges were activated with 1 mL acetonitrile, equilibrated with 1 mL water, loaded with samples, and washed with 1 mL water containing 1% formic acid. Peptides were eluted with 700 μL of 70% acetonitrile, dried under vacuum, and resuspended in 200 μL water. Peptide concentrations were determined using the Pierce BCA assay (Thermo Fisher Scientific, A55864) according to the manufacturer’s high-range NanoDrop protocol. Samples were adjusted to 1 mg/mL using the initial LC mobile phase (97% water, 2.9% acetonitrile, 0.1% formic acid).

### Mass Spectrometry-based Proteomic measurements

Peptides were separated using a Dionex Ultimate 3000 UHPLC from ThermoFisher in conjunction with an Acclaim PepMap 100 C18 HPLC column (150 x 0.3 mm, length x inner diameter; Part No. 164537). The separation was performed in the reverse-phase (RP) mode with 99.9% H_2_O and 0.1% FA (Solvent A), and 99.9% ACN and 0.1% FA (Solvent B). A 5 μL/min flow rate was maintained for all LC operations, and the analytical column was held at a constant temperature of 55 °C. Samples were injected via an autosampler fitted with a 20 µL sample loop and loaded to the analytical column at a rate of 5 µL/min. Additionally, sample injection order was randomized to mitigate the effect of instrumental drift on results. The autosampler was operated in the “Low Dispersion Mode” with a 1.7 low dispersion factor. The LC gradient consisted of a 6-min isocratic hold at 3% B, followed by a linear gradient from 3% to 35% B over 60 min, a 14-min wash at 95% B, and a 10-min re-equilibration at 3% B. Additionally, 1.5 μg BSA and 15 μg *E. coli* QCs were injected every 5 and 10 samples injections, respectively as per Bittremieux et al. (2018). No samples were excluded on the basis of QC performance.

Peptides were ionized using heat-assisted electrospray ionization (HESI) with a 2.5 kV spray voltage. Mass Spectrometric measurements were performed using a Q-Exactive HF-X from ThermoFisher Scientific. Data independent acquisition (DIA) was used to acquire peptide mass spectra. For all collected spectra the S-Lens RF level was set to 35. The capillary temperature was set to 250 °C, the sheath gas was set to 40, the aux gas was set to 3, and the probe heater was set to 25 °C. Importantly, a series of studies have explored the viability of low data point per peak (DPPP) DIA acquisition methods, and found acquisition rates as low as 1.5 and 1.25 DPPP to retain quantitative accuracy while improving proteomic coverage (Doellinger et al. 2020, 2023; Gu et al. 2024). In light of these findings, data was acquired using DIA mode with a resolving power of 30,000 using m/z 6.5 isolation window widths each offset by m/z 5.9407 with m/z 0.28 flanking overlaps spanning from m/z 400 to 1,000. As a result, 101 windows were established to scan from m/z 400-1000. Because these acquisitions occurred at 15 Hz across an estimated 15 s full width half max (FWHM) chromatographic peak width, the resultant DPPP can be estimated at 2.23. Fragment ion MS^2^ scan maximum injection time was set to 54 ms, automatic gain control (AGC) target was set to 3×10^6^, and higher-energy collisional dissociation (HCD) energy was set to 28.0. Full-scan spectra were collected once for every 100 DIA MS^2^ scans. Full scans were collected from m/z 400-1250 at 120,000 resolving power, with a 200 ms maximum ion injection time, and a 3×10^6^ AGC target.

### Proteomics data processing: Protein inferences and Quantitation

Raw DIA spectra were processed using MaxQuant (v2.6.7.0) (Cox and Mann 2008). Samples were analyzed without fractions, and the “type”option under “group-specific parameters” was set to MaxDIA with the predicted Library type and default fragmentation model type (Sinitcyn et al. 2021). The predicted library was formed using the *G. obscurus* (DSM 43160) reference proteome from Uniprot (UP000001382). Methionine oxidation and acetyl N-terms were included as variable modifications, and carbamidomethylation of cystine residues set as a fixed modification. LFQ was enabled with default settings, including a minimum ratio count of 2 (Cox et al. 2014). Trypsin/P was set for the digestion enzyme with 1 maximum missed cleavage. Minimum peptide length was set to 7 and maximum peptide mass was set to 4,600 Da.

### Bioinformatics Analysis

Bioinformatic analyses were performed using Perseus software (v2.1.3.0) (Tyanova et al. 2016). Uniprot accession numbers were annotated using the Perseus “Add annotation” function with the mainAnnot.bacteria.txt file (accessed at https://datashare.biochem.mpg.de/s/qe1IqcKbz2j2Ruf?dir=/PerseusAnnotation/Current/TaxonomicDivisions) (Boutet et al. 2007). Reverse sequences, contaminants, and proteins only identified by site were removed. LFQ intensities were log_2_ transformed, and proteins were retained only if they contained valid values in at least 70% of samples within one treatment group. Differential protein abundance between treatment groups was assessed using Perseus volcano plot analysis based on a two-sided Student’s t-test with 250 permutations and a 5% false discovery rate (FDR). Functional enrichment was assessed using the one-dimensional gene enrichment (1DGE) analysis implemented in Perseus, using mean log₂ fold changes between treatment groups as the ranking metric. Multiple hypothesis testing was corrected for among 1DGE results using Benjamini-Hochberg (BH) with an FDR cutoff of 5%. Figures were generated in R using ggplot2, displaying annotations exhibiting at least a 10% change in abundance.

### Data Deposition

The raw proteomic measurements (.RAW files) and MaxQuant analysis and corresponding output files have been deposited to the ProteomeXchange Consortium via the PRIDE partner repository under the accession number PXD086715.

## Results and Discussion

### UV Exposure Conditions and Proteome Coverage

Analysis of colony-forming unit (CFU) counts identified a median lethal UV dose (LD50) of approximately 39 mJ/cm², corresponding to a 13-s exposure under the experimental conditions used in this study (**Fig. 1**). Accordingly, all subsequent UV irradiation tests were performed using the same 13 seconds exposure. Proteomic analyses identified an average of 2,704 ± 38.7 SD protein groups were detected in each sample after contaminant sequence filtration was performed (**Fig. 2**). After filtering for proteins with valid values in at least 70% of samples within one treatment group, an average of 2,692 ± 36.3 SD protein groups remained per sample (**Fig. 2**). When comparing treatment groups, no statistically significant changes were observed at the protein level (**Fig 3**). Therefore, subsequent analyses focused on gene ontology (GO) enrichment rather than individual protein abundance (Ashburner et al. 2000). Although no individual proteins were significantly different in abundance between treatment groups, GO enrichment analysis identified functional categories exhibiting coordinated shifts in protein abundance, indicating that biologically meaningful changes occurred at the pathway level.

**Figure 1.**
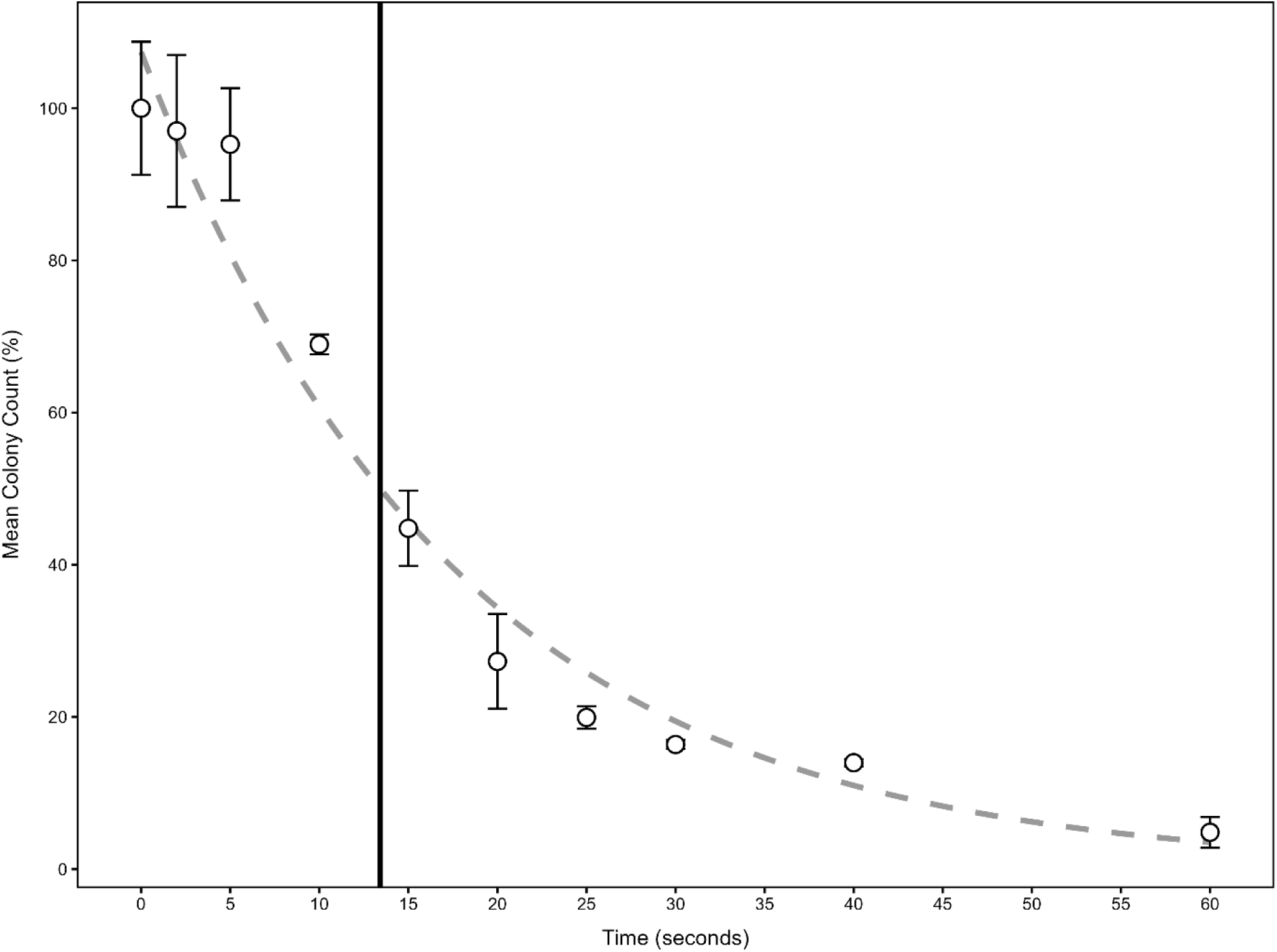
Viability of *Geodermatophilus obscurus* G20 under varied UV exposure times. G. obscurus streaked plates were each exposed to 254 nm UV irradiation for various durations ranging from 0 to 60 seconds. Resulting CFU counts were averaged by timepoint and normalized with respect to the CFU count at 0 seconds. An exponential decay line was fitted to the trend, and the point at which mean colony count was projected to be 50% was found to be at about 13 seconds, as denoted by the vertical line intersecting the trendline.

**Figure 2.**
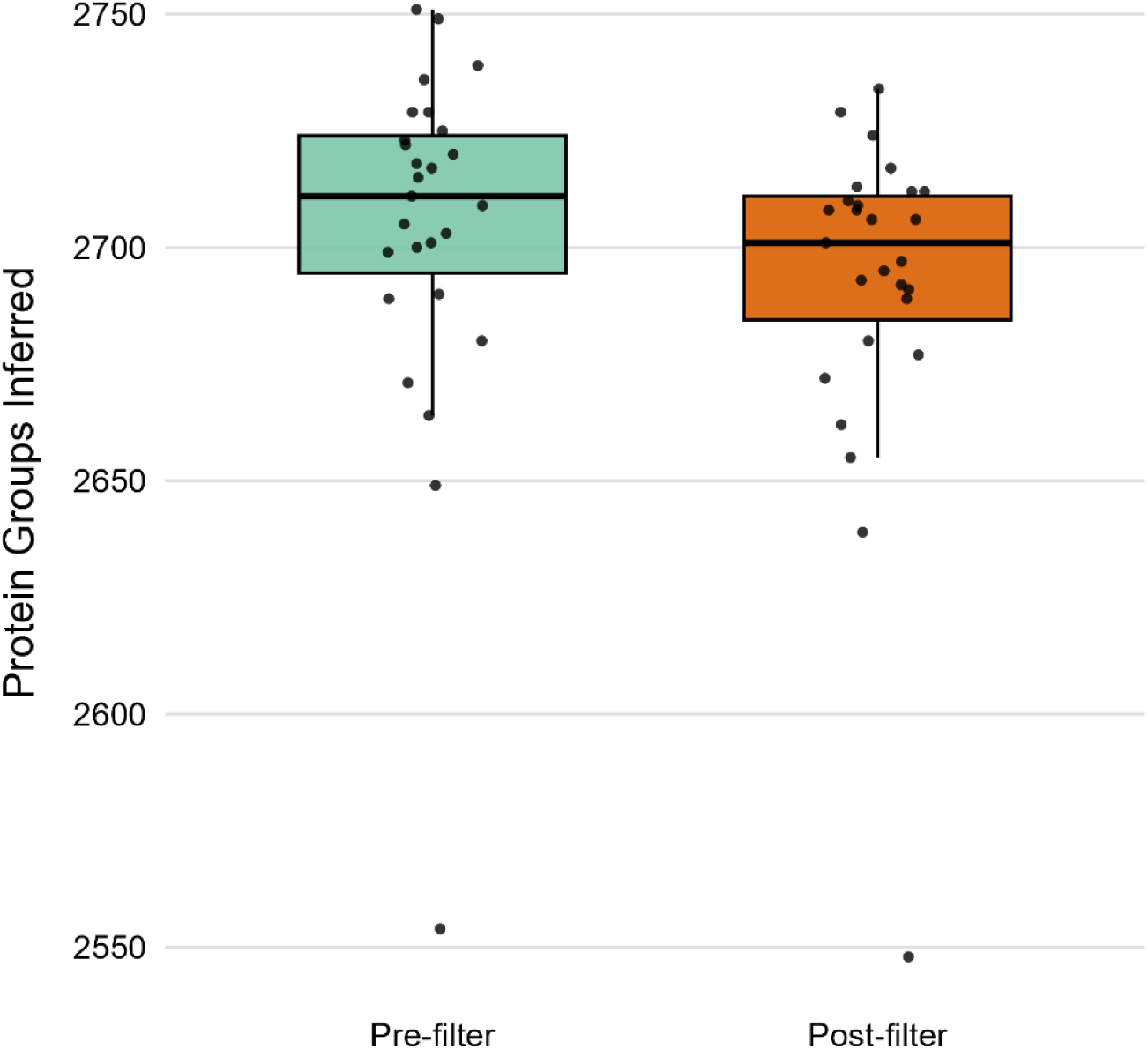
Protein Group Inferences Before and After Filtration. Total number of protein group inferences are depicted before and after valid-value filtration (n = 27). Pre-filter numbers are indicative of total LFQ quantifiable proteins observed in each sample. Post filter numbers are indicative of LFQ quantifiable protein groups which were observed in at least 70% of samples within at least one treatment group.

**Figure 3.**
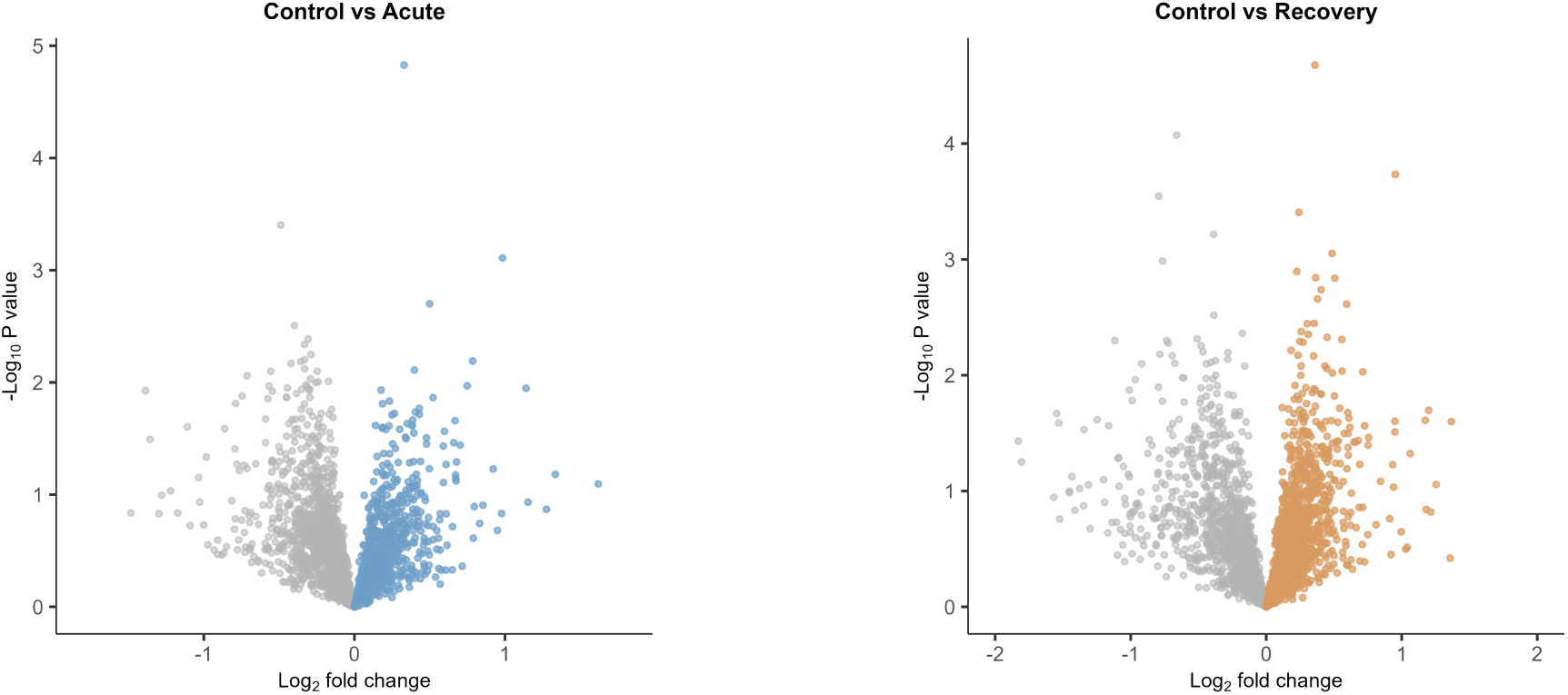
Quantitative Difference Among Protein Groups Between Treatment Groups. Comparisons made between treatment groups and control groups were carried out via student’s t-test with 250 permutations to correct for multiple hypothesis testing. In both acute (A) and recovery (B), no proteins were observed to be differentially abundant with statistical significance.

### Early UV response is characterized by increased NADH dehydrogenase activity

Changes in protein levels? between control and acute treatment groups were modest, and narrowly constrained to an increase in NADH dehydrogenase (NDH) activity and quinone binding (**Fig 4**). It is worth noting that these are automated annotations based on sequence homology, and *G. obscurus* G20 is known to specifically produce menaquinone (Collins et al. 1977; Montero-Calasanz et al. 2014). This suggests an increase in energetic demand, as NDH activity is largely responsible for the transfer of electrons from NADH to quinone to facilitate their introduction into the electron transport chain (ETC) (Sazanov 2015). There is a degree of ambiguity in this interpretation, however, as NDH is also responsible for redox homeostasis. Examination of the proteins contributing to this enrichment revealed that eight of the nine proteins corresponded to subunits of the nuo operon, with NuoL exhibiting the greatest increase in abundance (+44.8%) followed by NADH-quinone oxidoreductase (+36.2%) (Table S1). Specifically, subunits L, F, B, C, I, N, J, and D were identified in this matter. The dramatic increase in NuoL is also telling, as it is primarily involved in proton pumping as opposed to electron transfer (Nakamaru-Ogiso et al. 2010; Narayanan et al. 2016). This level of detail and NDH I specific contribution suggests that the primary form of contribution of these up-regulations is to facilitate a greater supply of energy to the afflicted cells. On the other hand, if redox homeostasis were the primary concern, Type II NDH activity would be the more likely observed response.

**Figure 4.**
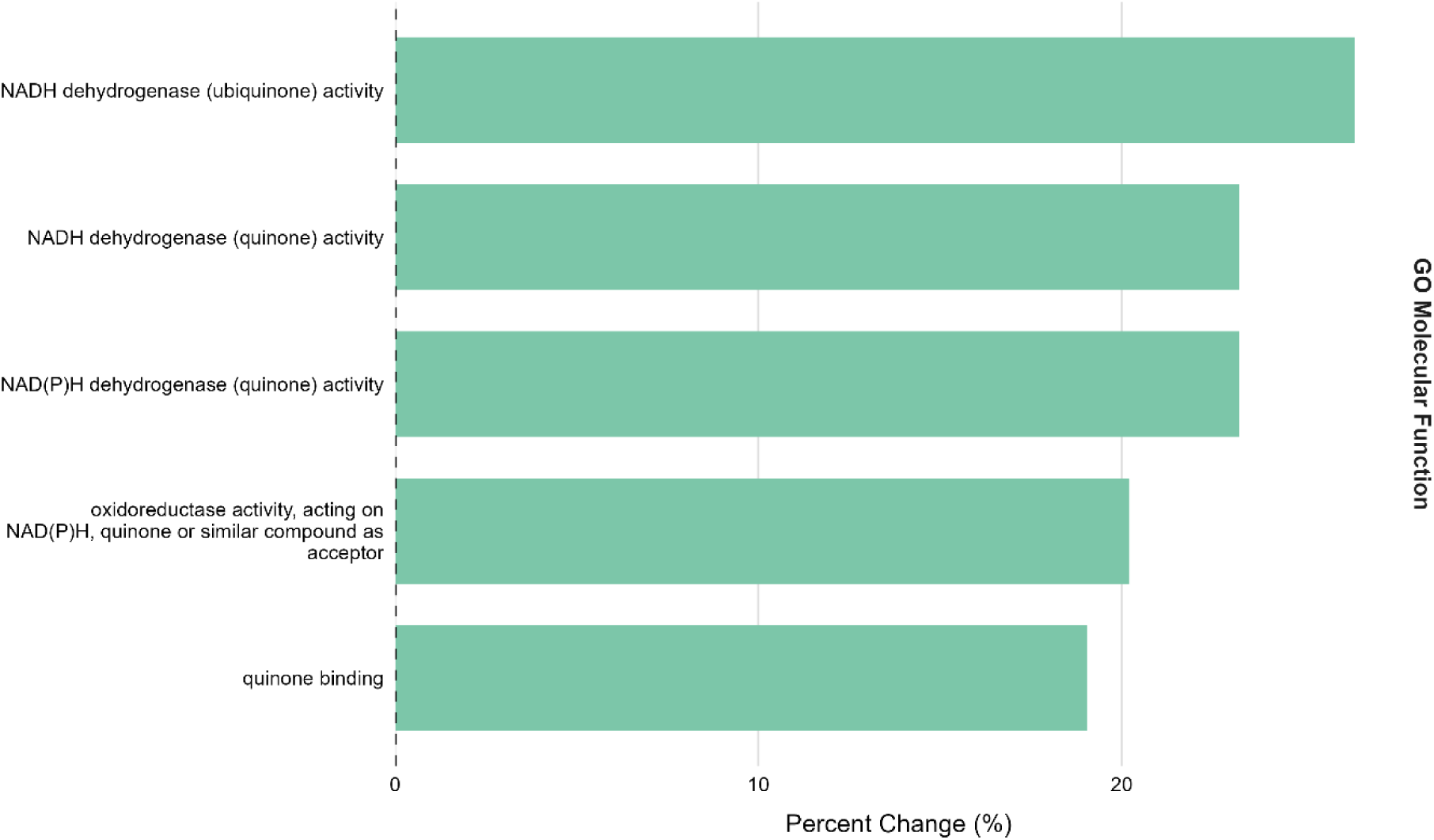
Differential enrichment of GO terms among acute treatment group samples. The mean difference in protein group intensity was calculated by subtracting the control group intensities from that of the acute treatment group. Subsequent 1DGE analysis revealed increases in NDH and quinone binding activity ranging from 19-26% with statistical significance (<5% FDR).

### Recovery phase reveals translational activation and reduced cell wall synthesis

Proteomic remodeling became more pronounced after the 1-h recovery period. Go enrichment analysis identified 87 significantly enriched annotations at a 5% FDR threshold (Table **S1**). Applying a minimum effect-size threshold of 10% reduced this to 13 annotations (**Fig 5**). Here, elevated NDH activity persisted during recovery, accompanied by additional changes associated with translation and cell wall synthesis. Annotations associated with translation (large ribosomal subunit and ribosomal subunit) were enriched, together with a 7.8% increase in the annotation “catalytic activity, acting on DNA” (Table S1), consistent with increased synthesis of proteins involved in DNA repair. Conversely, annotations associated with peptidoglycan and cell wall synthesis (carboxypeptidase binding, penicillin binding, and monocarboxylic acid binding) decreased (**Fig 5**), suggesting reduced cell wall biosynthesis (Macheboeuf et al. 2006; Sauvage et al. 2008). The largest individual decrease was observed for a peptidoglycan glycosyltransferase (−54.8%), which shared >90% sequence homology with the septal peptidoglycan synthetase FtsI. This finding is consistent with reduced septal peptidoglycan synthesis and, consequently, a transient reduction in cell division during recovery (Wissel and Weiss 2004). This indicates a down regulation of septal synthesis as well as cell division, which is reasonable to observe in an actively recovering cell (Chauhan et al. 2006). This supports a hypothesis that 1 hour post exposure cell division is slowed, and this stalling of cell division likely coincides with irradiation associated repair processes. Together, the persistence of elevated NDH activity and the apparent reduction in cell division suggest that recovering cells reallocate resources toward repair rather than growth. This interpretation is consistent with Patel et al. (2009), who reported transcriptional activation of growth-resumption pathways beginning approximately 30 min after UV exposure and peaking after 3 h. Accordingly, incomplete restoration of cell division after 1 h would be expected. Many of these changes were also observed when comparing recovery and acute treatment groups (Table S1) supporting the conclusion that they reflect biological recovery following UV exposure rather than sample-handling artifacts.

**Figure 5.**
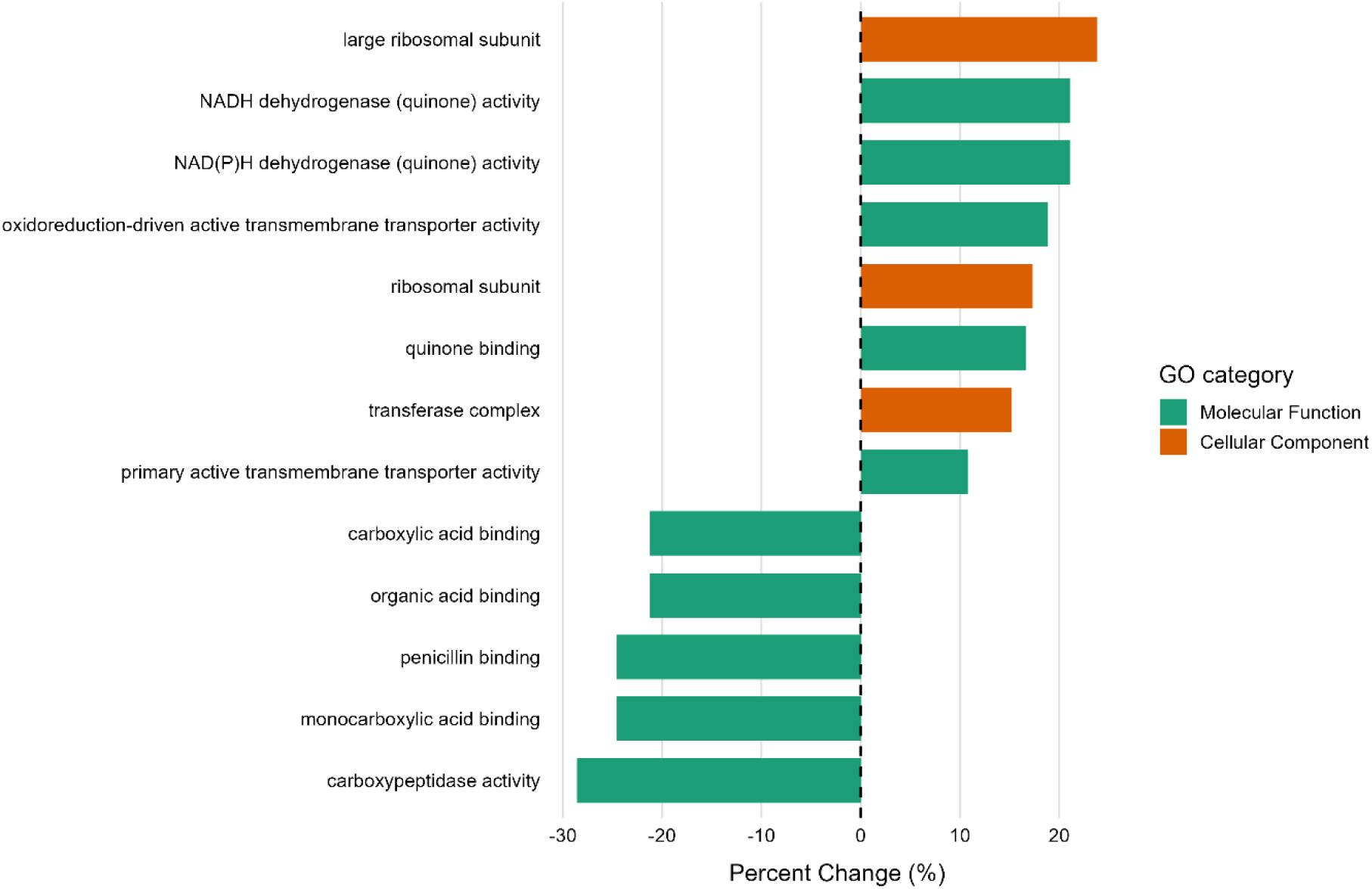
Differential enrichment of GO terms among recovery treatment group samples. The mean difference in protein group intensity was calculated by subtracting the control group intensities from that of the recovery treatment group. Subsequent 1DGE analysis revealed similar increases in NDH and quinone binding along with corresponding increases in translational activity, and decreases in cell wall synthesis with statistical significance (<5% FDR).

### Experimental limitations

Several limitations of the present study should be considered. The most important limitation is the clumped growth morphology of G. obscurus G20 and its potential effect on UV dose uniformity. Because *G. obscurus* G20 forms multicellular aggregates (Ivanova et al. 2010), cells located at the surface of a clump are likely to receive the full UV dose, whereas cells deeper within the aggregate may experience substantially attenuated irradiation due to a shadowing effect. In the current study, efforts were made to minimize aggregation by mechanically homogenizing starter cultures using a sterile pestle followed by repeated passage through a 26G syringe needle prior to inoculation. In addition, cell suspensions were vortexed immediately before UV irradiation to further disperse aggregates. Nevertheless, complete disruption of the characteristic multicellular clumps formed by *G. obscurus* G20 could not be confirmed. Consequently, heterogeneous UV exposure may have reduced the apparent magnitude of individual protein abundance changes by averaging highly irradiated and partially shielded cells within the same sample. This may partially explain the absence of statistically significant changes at the individual-protein level despite coordinated shifts detected by GO enrichment analysis. Combined with the UV absorption properties of its carotenoids (Sghaier et al. 2016) aggregate formation may therefore contribute substantially to the apparent UV resistance of this organism.. However, the demonstrated resistance of *G. obscurus* G20’s to mitomycin C (Gtari et al. 2012; Sghaier et al. 2016) indicates that resistance cannot be explained solely by UV shielding, since chemically induced DNA crosslinking is unaffected by carotenoid-mediated absorption. Selection of the UV dose also represents an important experimental consideration. Excessive irradiation may kill cells before a coordinated proteomic response can develop, whereas insufficient irradiation may fail to induce detectable physiological changes.

### Conclusions and future perspectives

This study provides the first proteomic characterization of the response of *Geodermatophilus obscurus* G20 to UV irradiation and early recovery. Although no individual proteins exhibited statistically significant changes in abundance, coordinated pathway-level responses revealed distinct physiological adaptations. Acute UV exposure was primarily associated with increased NADH dehydrogenase activity, while the recovery phase was characterized by sustained energy metabolism, increased translational activity, and reduced cell wall synthesis, consistent with a transient shift in cellular resources from growth toward recovery and repair.

Results from this study also suggest a focus for future experiments to improve the uniformity of UV exposure by further minimizing cell aggregation, and optimizing irradiation doses and recovery timepoints accordingly. Future studies could also compare UV-induced proteomic responses with those elicited by mitomycin C, a DNA crosslinking agent, to identify stress response pathways shared across distinct types of DNA damage. Such comparisons would require independently optimized treatment conditions to achieve comparable levels of cell survival.

Such studies will provide a more elaborate understanding of the molecular mechanisms underlying the remarkable UV resistance of *G. obscurus* G20 and other radiation-tolerant actinomycetes.

## Acknowledgements

The Microbial Ecology Collaborative funded this research, with funding from National Science Foundation (NSF) EPSCoR grant number #EPS-1655726.

## Data Availability Statement

The proteomic datasets generated in this study are publicly available on the ProteomeXchange Consortium via the PRIDE partner repository with a corresponding accession number of PXD086715.

